# Genetic diversity and connectivity of the invasive gastropod, *Callinina georgiana* (Caenogastropoda: Viviparidae) across a fragmented riverscape: a mitonuclear perspective

**DOI:** 10.1101/2024.02.24.581858

**Authors:** Wijesooriya Arachchilage Nimanthi Upeksha Abeyrathna, Andrew A. Davinack

## Abstract

1. Aquatic Invasive Species (AIS) are a significant threat to global freshwater biodiversity. This study focuses on the banded mystery snail, *Callinina georgiana,* an invasive species in the Adirondack region of northern New York – an important section of the New York Great Lakes Basin. This project aims to explore the genetic connectivity of *C. georgiana* within its invasive range using a combination of mitochondrial and nuclear markers.
2. Sampling was conducted in the Raquette River and adjacent waterways, with a total of 229 snails collected from 16 distinct populations distributed across eight different waterbodies. Also included were two populations from the species’ native range in the southern United States. DNA was extracted, and a 710 bp fragment of the mtDNA marker COI and a 351 bp fragment of nuclear marker Histone-3 (H3) were amplified. Population genetic analyses including haplotype patterning, AMOVA and genetic diversity estimates, neutrality tests and tests for isolation by distance (IBD) were performed to assess connectivity patterns.
3. Results showed moderate to high levels of genetic admixture within the snail’s invasive range as indicated by the lack of geographic patterning of haplotypes and low to moderate levels of genetic differentiation across multiple sites. Demographic analyses combined with high numbers of private haplotypes indicate population expansion. Interestingly, a case of mitonuclear discordance was detected for native and invasive populations as evident by incongruent haplotype patterns for the COI and H3 markers.
4. *Callinina georgiana* exhibits a high level of genetic connectivity in its invasive range. The presence of dams does not significantly affect gene flow, indicating that anthropogenic activities, such as boat traffic might be key in dispersing the snails across this fragmented freshwater system.
5. This study offers new insights into the dispersal and genetic structure of an invasive freshwater snail. It highlights the importance of considering anthropogenic factors when confronting complex patterns of genetic diversity. The findings are significant for biodiversity conservation and provide a basis for developing strategies to manage and contain the spread of AIS like *C. georgiana,* especially in regions with high human activity.

## 1. INTRODUCTION

Aquatic invasive species (AIS) pose one of the greatest threats to global freshwater biodiversity (Sala et al., 2000; Havel et al., 2015). The translocation of freshwater biota across wide geographic regions due to intentional (e.g., aquaculture stocking, ornamental trade) and unintentional (hitchhiking) vectors has resulted in the ‘dampening’ of biogeographic barriers and introduction of non-native species at a rate unprecedented in human history (David, 2018). While most introduced species never become invasive, those that do can severely disrupt communities through competition with native species, habitat modifications and even alterations of local biogeochemical cycles (Molnar et al., 2008, Mainka & Howard, 2010). In addition, freshwater systems provide important goods and services to human societies and therefore disruptions in their functioning incurs direct economic costs which are independent of those caused by biodiversity loss (Pimentel et al., 2005).

Of all the recent (∼30 years) freshwater invasions that have occurred around the world, some of the most high-profile invaders have been mollusks. For example, the zebra mussels (*Dreissena polymorpha polymorpha*), native to the Ponto-Caspian basin of Europe, has been the poster child for the negative impacts of invasive species, due to its role in both the abiotic and biotic transformation of the Great Lakes of North America, which some believe was one of the triggers for an invasion meltdown in this system (Ricciardi and MacIsaac 2000, but see also DeVanna et al., 2011). The spread of gastropods from Asia and Australasia to Europe and North America has garnered significant attention due to the speed at which they have spread across their new ranges. For example, the New Zealand mudsnail (*Potampyrgus antipodarum*) is now established on every continent except Antarctica and has been found in densities as high as 500,000 m^-2^, a feat that can probably be attributed to its reproductive mode, where all invasive populations are exclusively asexual (Alonso & Castro-Diez, 2012). Another example is the spread of the so-called ‘mystery snails’, native to Asian lakes and rivers, but invasive in many North American water bodies (Fowler et al., 2022). For example, the Japanese mystery snail, *Heterogen japonica* is widely distributed across North America where it thrives in the warm lakes of southern Texas but is also able to overwinter and spread in the water bodies of New England and Canada (David and Cote 2019). Meanwhile, other recent invaders such as *Sinotaia quadrata* and *Cipangopaludina* (=*Bellamya*) *chinensis*, both of which can harbor medically important parasites, have established themselves in a patchy distribution across most of the northern parts of the US (David & Cote, 2019; O’Leary et al., 2021).

Population genetic analyses have become standard protocol for understanding the spread and management of invasive species (Abdelkrim et al., 2005; Le Roux & Wieczorek 2009). This is because it provides a wealth of important data regarding the connectivity patterns of spatially separated populations, comparative genetic diversity estimates and demographic data regarding population expansions and contractions. Altogether, this information can be used to determine the long-term viability of an invader while also aiding in the development of management and or eradication strategies.

The banded mystery snail, *Callinina georgiania* (I. Lea, 1834), (formerly *Viviparus georgianus*) is a freshwater snail narrowly restricted to the eastern United States (Jokinen et al., 1982; Jokinen, 1992). The species is native to the south-eastern US and regarded as introduced in the northeastern and midwestern states (David et al., 2017). The earliest record of an introductory event for *C. georgiana* was in the mid-1800s when an aquarist released 200 specimens into the Hudson River. Since then, the snail has been introduced independently in other water bodies and has now successfully spread and colonized northern lakes and rivers including southernmost regions of Canada (Figure 1). *Callinina georgiana* is rare among gastropods in that it is ovoviviparous, producing 4-80 live young with limited dispersal capabilities (Jokinen et al., 1982; Jokinen, 1992). The snail is a species of concern for environmental authorities in states like New York, Minnesota and Michigan which border the Great Lakes basin, where its impacts have been most felt. *Callinina georgiana* is also an egg predator of largemouth bass and reaches high densities in small ponds and lakes, outnumbering native macrofauna in the invaded area (Eckblad & Shealy Jr, 1972). Like many mystery snails, *C. georgiana* is also an intermediate host to medically important trematodes (David et al., 2017), which were implicated in a mass die-off of aquatic birds in 2007 on a lake in northern Wisconsin. Despite a comprehensive knowledge of *C. georgiana’s* life-history, nothing is known about its actual dispersal capabilities nor its connectivity patterns in its native or invasive range.

**Figure 1.**
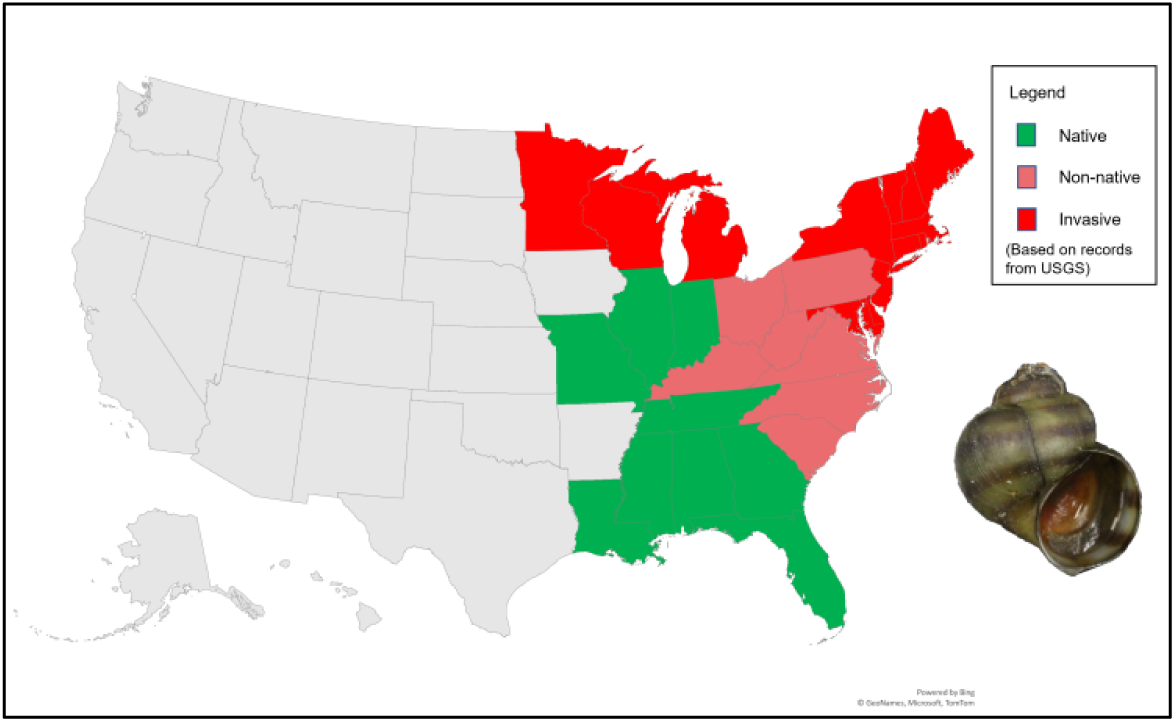
Distributional map of *Callinina georgiana* in the continental United States. The native range, non-native range and invasive range was defined according to the latest United States Geological Survey record (date accessed: 09/21/2022, date last updated: 03/20/2024

In this study, we aimed to explore the genetic connectivity of *C. georgiana* within its invasive range in the New York Great Lakes Basin (NYGLB), a significant portion of its non-native habitat. Our focus was on examining the snail’s dispersal ability in the fragmented Raquette River region and adjacent waterways. We utilized both mitochondrial and nuclear markers for this assessment. The Raquette River, New York’s third longest river, featuring 27 hydroelectric dams, which has been demonstrated in previous studies to disrupt hydrological connectivity for migratory species (Underwood et al., 2016; Zarri et al., 2022). Additionally, we included samples from two native populations in the south-eastern US in our analysis. Such inclusion is vital in invasion genetics to understand the dynamics of biological invasions, including invasion pathways, the genetic composition of initial populations, and the significance of genetic diversity in invasion success (Allendorf & Lundquist, 2003; Lawson Handley et al., 2011). This approach not only aids in identifying the transition point from eradication to containment efforts but also enhances risk assessment of invasive species (Stepien et al., 2005). Our goal was to use these genetic markers to (i) assess genetic structure of *C. georgiana* populations within the Raquette River, surrounding waterways, and a portion of its native range and (ii) evaluate genetic diviersity and and demographic trends, in these areas.

## 2. METHODS

### 2.1 Description of Study Region

The Adirondack region is home to the Adirondack Park, the largest protected forest reserve in the contiguous United States (6.1 million acres) and contains more than 10,000 lakes and 30,000 miles of rivers and streams. Of these water bodies, the Raquette River is by far the most important in terms of energy production and recreation in the region. At 146 miles in length, it is the third longest river in New York State and one of most dammed rivers east of the Mississippi basin. These dams have resulted in several man-made reservoirs which for decades have been a key driver of tourism in the region where recreational boaters in the summer frequently traverse the waterways and anglers take advantage of these artificially stocked ‘lakes’. The Raquette River commences in Blue Mountain Lake in the heart of the Adirondack mountains and extends northwards until it reaches a confluence with the larger St. Lawrence River which receives discharge from Lake Ontario to the west and empties into the Atlantic Ocean to the east. Along with other smaller sister rivers such as the Grasse River and the Oswegatchie River, this region is part of the larger New York Great Lakes Basin (NYGLB). For the southern sampling region, both Wheeler branch and Cypress creek represent two tributaries on the Tennessee River. The river stretches from the eastern edge of the State of Tennessee and crosses into Alabama, terminating near the border of Mississippi.

### 2.2 Collection and sampling design

Historical distributional records of *C. georgiana* in New York State compiled by Jokinen (1992) along with occurrence data from online platforms such as ‘iNaturalist’ and ‘iMapInvasives’ were used to select sampling sites for this study. Fieldwork was completed over four months (May to August 2021) at 28 sites, the majority of which were located on the Raquette River with additional sites on sister rivers such as the Grass River, Oswegatchie River in addition to Lake Ontario and isolated lakes within the Adirondack Park itself. A timed survey was conducted at each sampling site along the river banks for 20-30 minutes. Adult snails from each location were initially identified in the field based on the characteristic chestnut-colored bands on the shell’s exterior, and were collected by handpicking. Water quality data for each sampling site (temperature and pH) were also recorded (Supplementary Table 1). In total, 229 snails were collected representing 16 distinct populations and from eight different water bodies (Figure 2, Table 1, Supplementary Table 1, Supplementary Table 2). In addition, 24 individuals from the species’ native southern range were supplied by colleagues working in the southern United States. These snails were collected from two sites on the Tennessee River in Alabama (Table 1). Collected specimens were fixed in 95% ethanol in labeled sampling bottles and transported to Clarkson University for further processing. In the lab, snails were conchologically identified first using the taxonomic key of Jokinen (1992), and then delineated further with updated characteristics described in David et al. (2017). All sampling, handling and processing of specimens were approved by the New York State Department of Environmental Conservation (Collections Permit numbers: 6-21-001 & 6-21-002).

**Figure 2.**
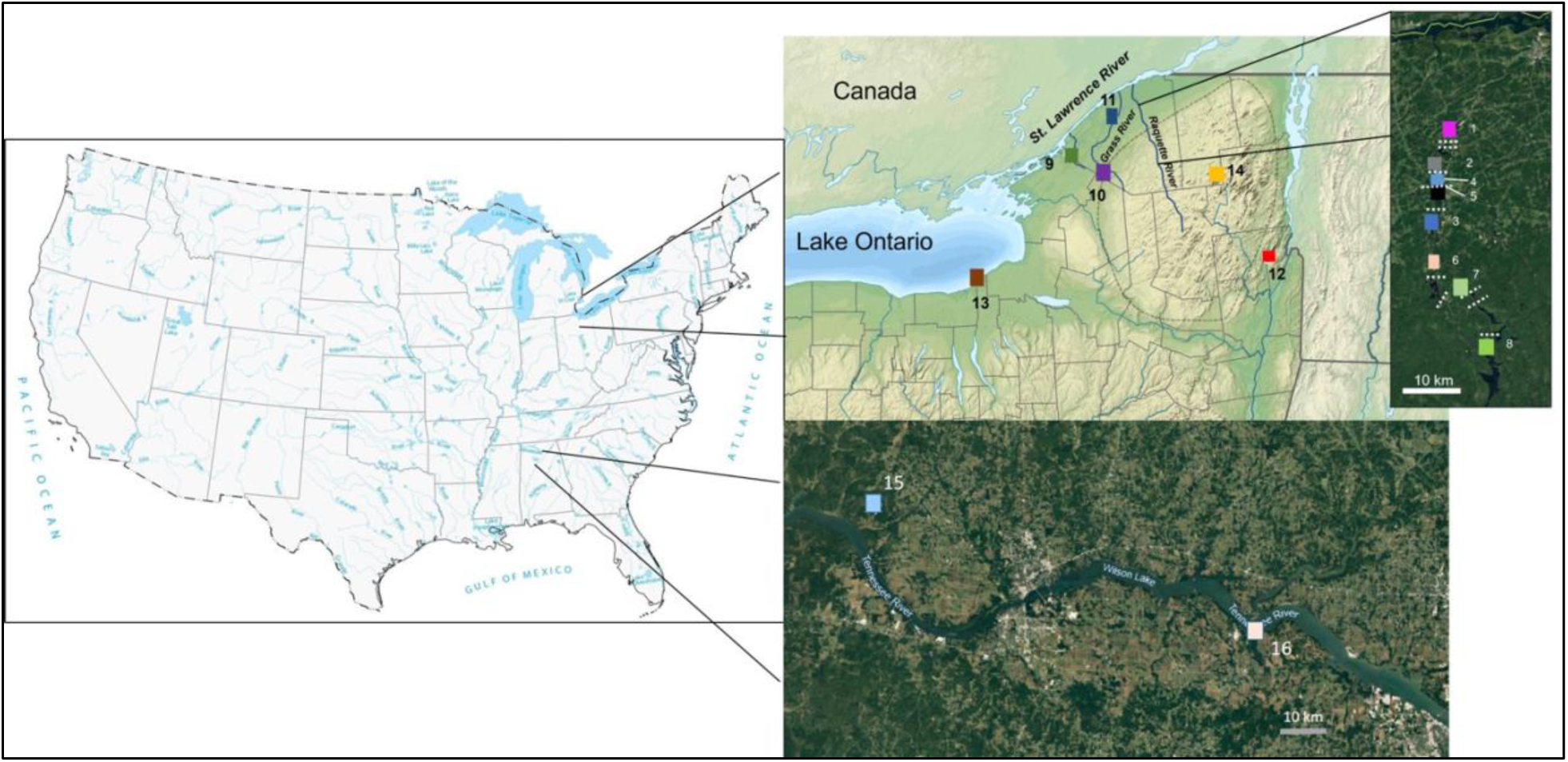
Sampling sites for *Callinina georgiana* snails collected from 16 distinct populations from six waterbodies, throughout the New York Great Lakes Basin. White dotted lines represent hydroelectrical dams present on the Raquette River (populations from 1-16 represents, Norwood, Sissonville, Hannawa, Potsdam 1, Potsdam2, Colton, South Colton, Stark Falls, Ogdensberg, Canton,Madrid, Lake George, Sodus Point, Paul Smiths, Cypress Creek and Wheeler Creek respectively).

**Table 1:**
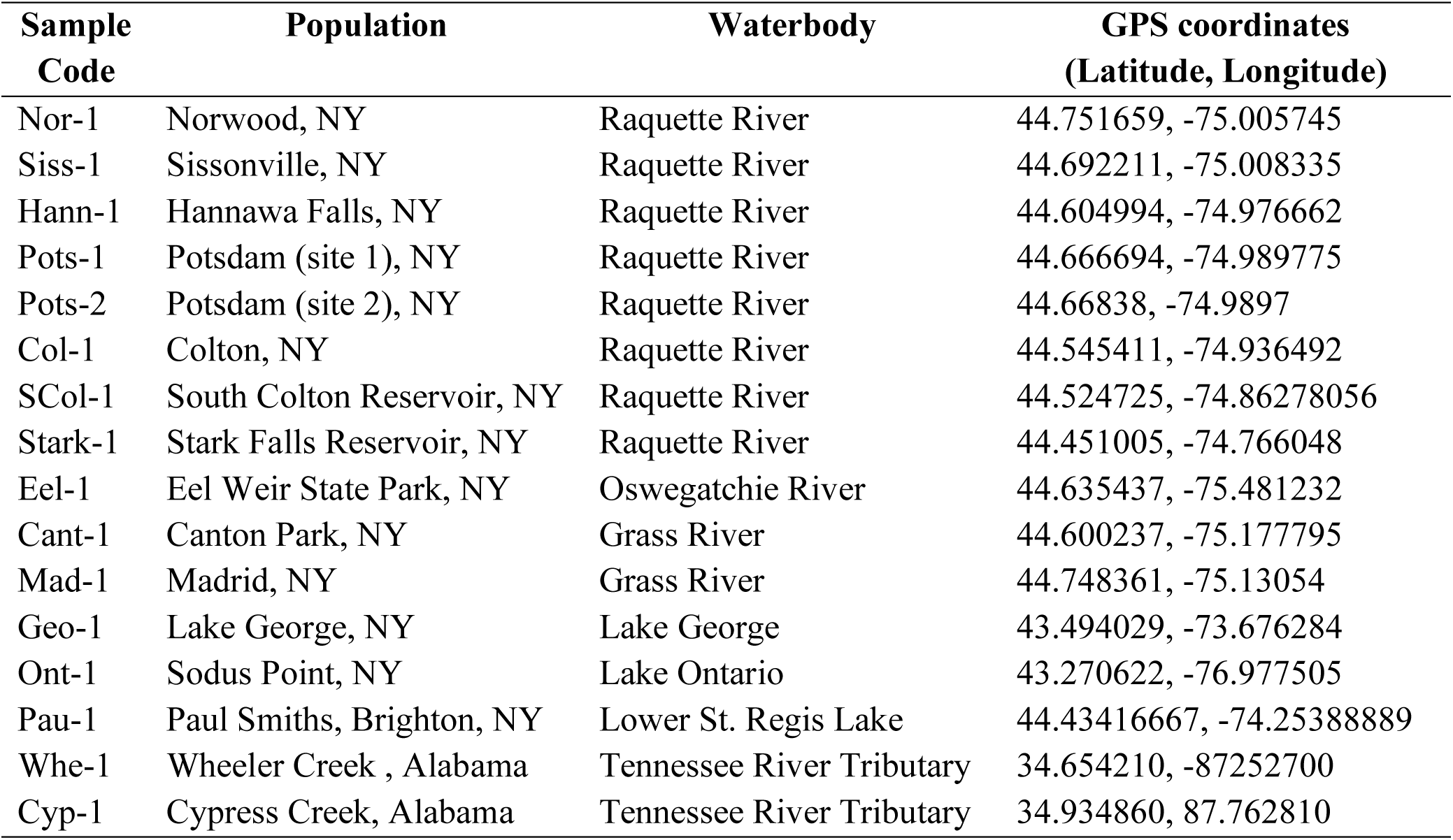
Upstate New York sampling locations for *Callinina georgiana*.

### 2.3 DNA Extraction and PCR Amplification protocol

For each population, a maximum of 10 snails per population was chosen for genotyping. Shells of specimens were cracked and a 0.25 mm section of the head tissue was digested in a Proteinase K and lysis buffer solution. Genomic DNA was extracted from a total of 172 snails using a D’Neasy DNA extraction kit (QIAGEN®, Hilden, Germany) with modifications to the manufacturer’s instructions (see David et al., 2017 and David & Cote, 2019). A 710 bp fragment of the mtDNA gene, cytochrome c oxidase 1 (CO1) and a 351 bp fragment of nuclear gene marker, Histone-3 (H3) were amplified using the universal forward and reverse primer pairs of Folmer et al., (1994) (LCO1490 and HCO2198) and Hayes et al. (2008) (H3F and H3R). Polymerase Chain Reaction was carried out in a 25 µl reaction mixture with the following cycling conditions for COI: initial denaturation of 95°C for 5 mins, followed by 45 cycles of 95°C for 1 min, an annealing temperature of 50°C for 1 min, 72°C for 1 min and a final extension of 72°C for 10 mins. H3 cycling conditions followed that of Hayes et al. (2008) with annealing temperatures of 55°C and 58°C. All amplicons were visualized on a 1% agarose gel and sequenced using both the forward and reverse primers at Azenta LLC (South Plainfield, NJ). Similarity indices for each sequence was checked using the BLASTn tool in GenBank and confirmed the sequences belong to *C. georgiana* with higher percentage identities. Finally, the sequences were checked for gene functionality using the ExPASY translation tool. All sequence data were deposited into GenBank (accession numbers: OP546054 – OP546057, OP547442 – OP547448, OP548039 – OP548049, OP548083 – OP548097, OP548123 – OP548133, OP549713 – OP549747, PP346073- PP346093, PP375058 – PP375081).

### 2.4 Sequence alignment, editing and population genetic analyses

Compiled sequences were aligned and edited using the ClustalW alignment tool in BioEdit ver.7.2 (Hall, 1999). After editing, a 559 bp COI fragment remained for analysis without gaps or missing data, and a 305 bp fragment remained for the H3 marker. Overall, a total of 118 COI sequences and 141 H3 sequences were used for population genetic analyses. To determine evolutionary relationships and diversity among haplotypes, haplotype networks for both markers were constructed using a TCS network implemented in POPART ver 1.7 (Leigh and Bryant 2015). A hierarchical analysis of molecular variance (AMOVA) along with pairwise F_ST_ and Φ_ST_ calculations were carried out in Arlequin ver. 3.5.2 (Excoffier & Lischer, 2010) to determine the extent of genetic differentiation among *C. georgiana* populations. Genetic diversity estimates for both markers were also calculated, first for the native and invasive range and then for each population along with neutrality tests (Fu’s Fs and Tajima’s D) in DnaSP ver. 6.0 (Rozas et al., 2017). Finally, a test for isolation by distance (IBD) using Mantel Test (Telles & Diniz Filho, 2005; Bohonak, 2002) to infer the extent that geographic distance influenced genetic distance was carried out using R packages geodist, ape and vegan. We categorized the sites with dams in the Raquette River as impounded sites (Nor1, Sis1, Han1, Potsdam 1, Potsdam 2, Colton 1, SCol-1, Stark 1) and other sites from different water bodies as unimpounded sites (Eel-1, Cant-1, Mad-1, Ont-1, Geo-1, Pau-1, Cyp-1, Whe-1). Isolation by distance is not expected on impounded sites due to multiple factors such barriers to gene flow, kin-structures migration, specific environmental factors such as hydrological parameters (Xiang et al., 2019; Duforet-Frebourg & Blum, 2014; Fix, 1993) and it may exhibit non-standard patterns of isolation by distance.

## 3. RESULTS

An analysis of the mtDNA COI marker recovered 80 haplotypes (10 shared and 70 unique) which exhibited marked geographic patterning with respects to the snail’s native and invasive range, but high levels of admixture was observed within the invasive range (Figure 3) The most common mtDNA haplotype was shared among individuals from six populations from the invasive range with the most distant populations being ∼107 km apart. A hierarchical analysis of molecular variance (AMOVA) showed that the majority of the genetic variation was found within populations (65.70%) rather than among them (34.30%) with a moderate level of structure that was significant (Φ_ST_ = 0.343, P < 0.01). Overall, pairwise Φ_ST_ ranged from 0.000 - 0.780 (Table 2) with the most structure found between the Sissonville (Siss-1) population on the Raquette River and the Wheeler Creek (Whe-1) population from a tributary in the Tennessee River (Φ_ST_ =0.780, P<0.05). The weakest structure (highest levels of connectivity) observed was between Sissonville (Siss-1) and Potsdam (Pots-1) (Φ_ST_=0.012, P>0.05). In addition, a Mantel test for impounded sites at the Raquette River did not find a statistically significant correlation between genetic and geographic distances (r=0.043, P=0.215) while results from unimpounded site showed a significant but weak correlation between geographic and genetic distance (r = 0.394, P= 0.001). The highest haplotype diversity (*h*=1.000) was found in Norwood, Potsdam 2, Hannawa and Stark Falls populations in the Raquette River and in Lake Ontario while the lowest reported was Madrid population (0.500) on the Grasse River. Overall, *C. georgiana* exhibited high haplotype diversity (*h* = 0.975 ± 0.008) and low nucleotide diversity (π = 0.017± 0.008) with populations from the native range exhibiting marginally higher haplotype diversity than populations from the invasive range (native *h* = 0.978; invasive *h*: 0.964). Demographic analyses of the species as a whole, yielded negative values (Fu’s F_s_= -2.535, P<0.05 and Tajima D=-1.728, P>0.05) (Table 3).

**Figure 3.**
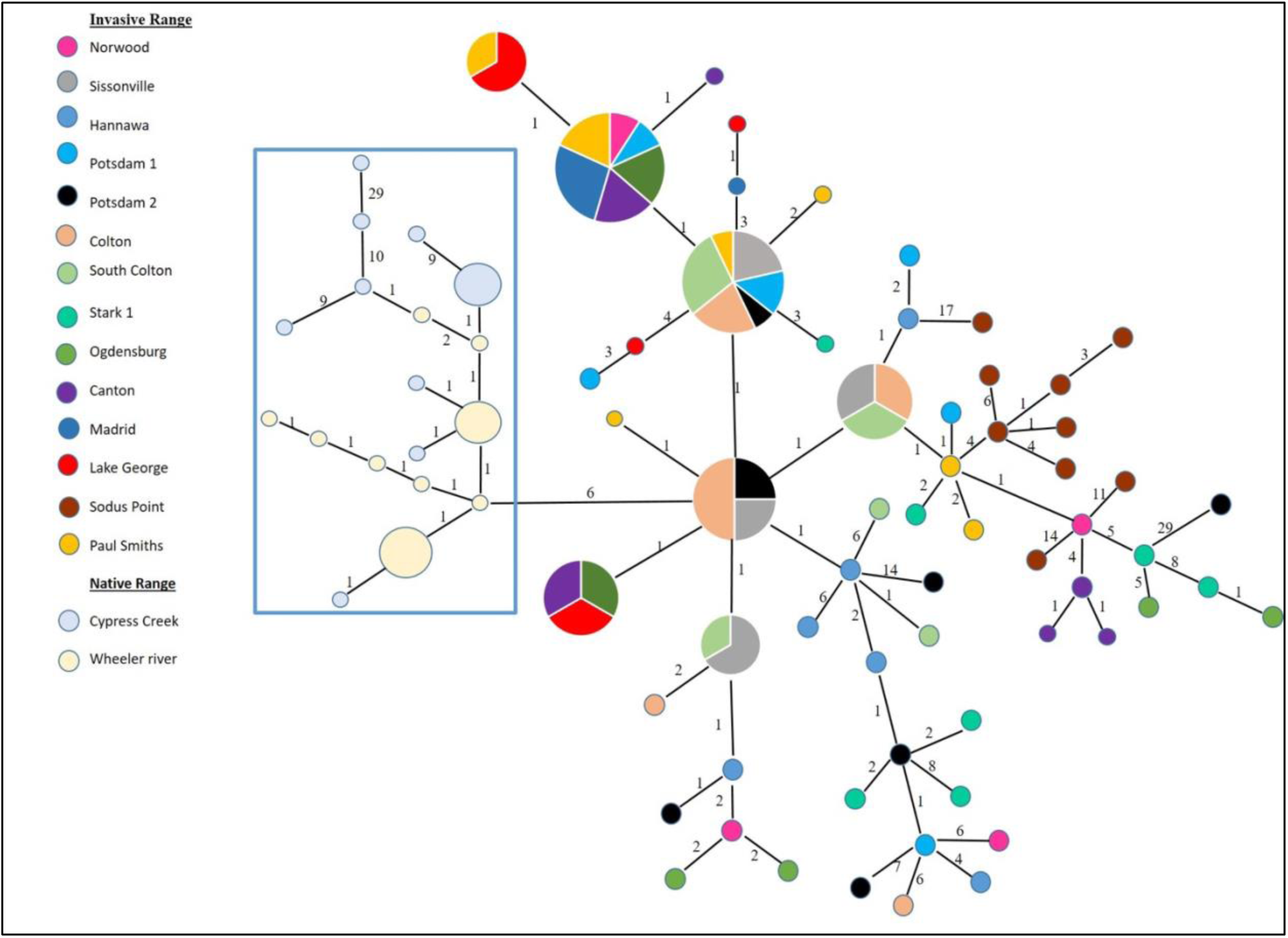
Haplotype network for *Callinina georgiana* based on mtDNA – COI sequence data. Size of circles is representative of the number of individuals with that haplotype. Smallest circles represent a haplotype frequency of one. Each connecting line between haplotypes represents one mutational step and numbers above lines represent the number of additional mutational steps.

**Table 2.** Pairwise genetic differentiation (Φ_ST_) among populations of *Callinina georgiana* based on the mtDNA COI data set. Bolded values denote significance (P < 0.05).

|  | Nor1 | Siss1 | Hann<br>1 | Pots1 | Pots2 | Col1 | Scol1 | Stark<br>1 | Eel1 | Cant1 | Mad1 | Geo1 | Ont1 | Pau1 | Whe1 | Cyp1 |
| --- | --- | --- | --- | --- | --- | --- | --- | --- | --- | --- | --- | --- | --- | --- | --- | --- |
| Nor1 | 0.000 |  |  |  |  |  |  |  |  |  |  |  |  |  |  |  |
| Siss1 | 0.087 | 0.000 |  |  |  |  |  |  |  |  |  |  |  |  |  |  |
| Hann1 | 0.048 | **0.251** | 0.000 |  |  |  |  |  |  |  |  |  |  |  |  |  |
| Pots1 | 0.030 | 0.012 | **0.144** | 0.000 |  |  |  |  |  |  |  |  |  |  |  |  |
| Pots2 | 0.072 | **0.119** | 0.025 | 0.068 | 0.000 |  |  |  |  |  |  |  |  |  |  |  |
| Col1 | 0.042 | 0.066 | 0.086 | 0.019 | 0.058 | 0.000 |  |  |  |  |  |  |  |  |  |  |
| Scol1 | 0.016 | 0.076 | **0.140** | 0.073 | 0.077 | 0.063 | 0.000 |  |  |  |  |  |  |  |  |  |
| Stark1 | 0.015 | **0.264** | 0.046 | **0.154** | 0.040 | **0.156** | **0.173** | 0.000 |  |  |  |  |  |  |  |  |
| Eel1 | 0.035 | **0.222** | **0.212** | 0.066 | 0.035 | 0.122 | **0.106** | **0.118** | 0.000 |  |  |  |  |  |  |  |
| Cant1 | 0.034 | **0.263** | **0.230** | **0.148** | 0.068 | **0.167** | **0.158** | **0.146** | 0.015 | 0.000 |  |  |  |  |  |  |
| Mad1 | 0.233 | **0.492** | 0.449 | 0.117 | 0.131 | **0.286** | **0.219** | **0.302** | 0.053 | 0.185 | 0.000 |  |  |  |  |  |
| Geo1 | **0.121** | **0.253** | **0.311** | 0.030 | **0.119** | **0.203** | **0.112** | **0.209** | 0.026 | 0.102 | 0.029 | 0.000 |  |  |  |  |
| Ont-1 | **0.176** | **0.299** | **0.269** | **0.257** | **0.213** | **0.278** | **0.275** | **0.298** | **0.323** | **0.184** | **0.341** | **0.342** | 0.000 |  |  |  |
| Pau1 | 0.131 | 0.111 | **0.349** | 0.036 | **0.156** | 0.100 | 0.029 | **0.283** | **0.083** | **0.332** | 0.073 | 0.010 | **0.330** | 0.000 |  |  |
| Whe1 | **0.668** | **0.738** | **0.691** | **0.650** | **0.506** | **0.662** | **0.673** | **0.630** | **0.669** | **0.705** | **0.780** | **0.683** | **0.615** | 0.710 | 0.000 |  |
| Cyp1 | **0.358** | **0.472** | **0.422** | **0.432** | **0.326** | **0.439** | **0.450** | **0.381** | **0.403** | **0.444** | **0.444** | **0.444** | **0.481** | 0.476 | 0.200 | 0.000 |

**Table 3.** Mitochondrial (CO1) gene polymorphism, genetic diversity and demographics analysis in *Callinina georgiana*. *N*= number of sequenced samples, *S*= number of segregating sites, *h*=haplotype diversity, *π*= nucleotide diversity.

| Population | *N* | *S* | *h* | $\pi$ | Fu's *F<sub>s</sub>* | Tajima *D* |
| --- | --- | --- | --- | --- | --- | --- |
| Nor-1 | 4 | 11 | 1.000 | 0.012 | 0.307 | 0.108 |
| Siss-1 | 7 | 3 | 0.809 | 0.002 | 0.741 | 0.796 |
| Hann-1 | 6 | 15 | 1.000 | 0.011 | -0.764 | -0.653 |
| Pots-1 | 7 | 16 | 0.952 | 0.010 | -1.154 | -1.132 |
| Pots-2 | 7 | 48 | 1.000 | 0.032 | -0.904 | -0.848 |
| Col-1 | 8 | 13 | 0.857 | 0.007 | -1.133 | -0.978 |
| SCol-1 | 8 | 13 | 0.786 | 0.007 | -1.296 | -1.016 |
| Stark-1 | 7 | 23 | 1.000 | 0.017 | -0.621 | -0.673 |
| Eel-1 | 6 | 14 | 0.933 | 0.009 | -1.440 | -1.359 |
| Mad-1 | 4 | 1 | 0.500 | 0.001 | -0.478 | -0.612 |
| Cant-1 | 7 | 7 | 0.952 | 0.006 | 0.824 | 0.875 |
| Geo-1 | 7 | 13 | 0.952 | 0.009 | -0.266 | -0.365 |
| Ont-1 | 10 | 40 | 1.000 | 0.022 | -0.703 | -0.716 |
| Pau-1 | 8 | 9 | 0.964 | 0.005 | -0.987 | -0.916 |
| Cyp-1 | 10 | 46 | 0.977 | 0.025 | -1.218 | -1.059 |
| Whe-1 | 12 | 6 | 0.939 | 0.041 | 0.353 | -0.019 |

For the H3 marker, a total of 58 haplotypes (24 shared and 34 unique) were recovered but unlike the COI data set, the H3 marker did not exhibit any marked geographic patterning of native and invasive haplotypes (Figure 4). The most common nuclear haplotype was shared among individuals from 10 populations distributed across both native and invasive ranges. A hierarchical AMOVA showed that the majority of the genetic variation was found within individual populations (64.17%) rather than among them (35.82%) with a moderate level of structure that was significant (F_ST_ = 0.358, P < 0.01). However, pairwise Φ_ST_ calculations did show higher levels genetic structuring among certain populations with the most structure observed between Potsdam1 (Pots-1) on Raquette River and Cypress creek (Cyp-1) on a tributary of the Tennessee River (Φ_ST_ = 0.903, P < 0.05), while the weakest structure was between Hannawa (Hann-1) and Colton (Col- 1) on the Raquette River and Potsdam 2 (Pots-2) on the Raquette River and Canton Park (Cant- 1) on the Grass River (Φ_ST_ = 0.017, P >0.05) (Table 4). Similar to the COI marker, a Mantel test for impounded sites at the Raquette River did not find a statistically significant correlation between genetic and geographic distances (r=-0.069, P=0.836). However, there was also no significant correlation for unimpounded sites (r =- 0.088, P= 0.99). Similar to the COI marker, *C. georgiana* exhibited high haplotype diversity (*h* = 0.975± 0.008) and low nucleotide diversity (π = 0.017) for the H3 marker. However, native populations exhibited comparably lower haplotype diversity (native *h*: 0.228) compared to invasive populations (invasive *h*: 0.960). The Norwood population (Nor-1) on the Raquette River had the highest haplotype diversity (1.000) while the Cypress population (Cyp-1) from the tributary of the Tennessee River had the lowest (0.000), the latter of which had no polymorphic sites. Similar to the COI data, demographic metrics were all negative (Fu’s Fs = -28.168, P < 0.05; Tajima D = -1.121, P > 0.10) (Table 5).

**Figure 4.**
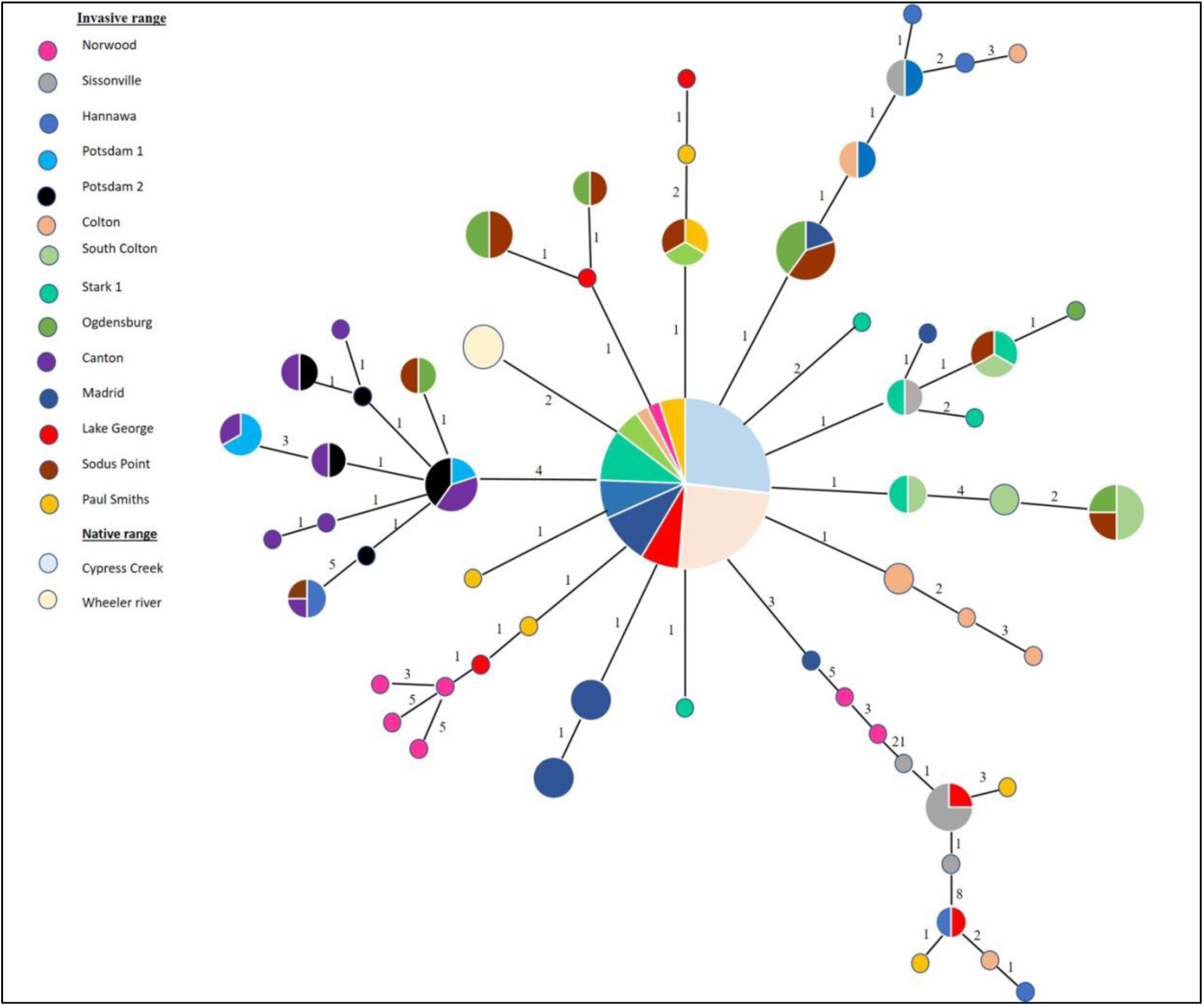
Haplotype network for *Callinina georgiana* based on nDNA – H3 sequence data. Size of circles is representative of the number of individuals with that haplotype. Smallest circles represent a haplotype frequency of one. Each connecting line between haplotypes represents one mutational step and numbers above lines represent additional mutational steps.

**Table 4:** Pairwise genetic differentiation (Φ_ST_) among populations of *Callinina georgiana* based on nuclear H-3 sequence data. Bolded values denote significance (P < 0.05)

|  | Nor1 | Siss1 | Hann1 | Pots1 | Pots2 | Col1 | Scol1 | Stark1 | Eel1 | Cant1 | Mad1 | Geo1 | Ont1 | Pau-1 | Whe1 | Cyp1 |
| --- | --- | --- | --- | --- | --- | --- | --- | --- | --- | --- | --- | --- | --- | --- | --- | --- |
| Nor1 | 0.000 |  |  |  |  |  |  |  |  |  |  |  |  |  |  |  |
| Siss1 | **0.545** | 0.000 |  |  |  |  |  |  |  |  |  |  |  |  |  |  |
| Hann1 | 0.130 | **0.468** | 0.000 |  |  |  |  |  |  |  |  |  |  |  |  |  |
| Pots1 | **0.514** | **0.613** | **0.358** | 0.000 |  |  |  |  |  |  |  |  |  |  |  |  |
| Pots2 | **0.602** | **0.688** | **0.390** | 0.225 | 0.000 |  |  |  |  |  |  |  |  |  |  |  |
| Col1 | **0.163** | **0.412** | 0.017 | **0.347** | **0.397** | 0.000 |  |  |  |  |  |  |  |  |  |  |
| Scol1 | **0.276** | **0.652** | **0.229** | **0.596** | **0.686** | **0.255** | 0.000 |  |  |  |  |  |  |  |  |  |
| Stark1 | **0.296** | **0.684** | **0.084** | **0.634** | **0.734** | **0.170** | **0.330** | 0.000 |  |  |  |  |  |  |  |  |
| Eel1 | **0.221** | **0.613** | **0.090** | **0.468** | **0.524** | **0.136** | **0.208** | 0.125 | 0.000 |  |  |  |  |  |  |  |
| Cant1 | **0.551** | **0.682** | **0.387** | 0.087 | 0.017 | **0.398** | **0.613** | **0.638** | **0.482** | 0.000 |  |  |  |  |  |  |
| Mad1 | **0.315** | **0.668** | **0.115** | **0.619** | **0.719** | **0.184** | **0.386** | **0.202** | **0.180** | **0.626** | 0.000 |  |  |  |  |  |
| Geo1 | 0.122 | **0.272** | 0.019 | **0.328** | **0.374** | 0.033 | **0.262** | **0.193** | 0.157 | **0.381** | **0.198** | 0.000 |  |  |  |  |
| Ont1 | **0.202** | **0.594** | 0.081 | **0.368** | **0.420** | **0.129** | **0.200** | 0.114 | 0.035 | **0.388** | **0.160** | 0.146 | 0.000 |  |  |  |
| Paul1 | 0.132 | **0.285** | 0.039 | **0.344** | **0.399** | 0.051 | **0.257** | **0.204** | **0.168** | **0.404** | **0.208** | 0.040 | **0.160** | 0.000 |  |  |
| Whe1 | **0.375** | **0.719** | **0.109** | **0.709** | **0.817** | **0.213** | **0.469** | **0.171** | **0.223** | **0.700** | **0.265** | **0.213** | **0.194** | **0.250** | 0.000 |  |
| Cyp1 | **0.415** | **0.745** | **0.113** | **0.762** | **0.903** | **0.228** | **0.522** | **0.156** | **0.230** | **0.748** | **0.299** | **0.237** | **0.193** | **0.226** | **0.217** | 0.000 |

**Table 5.** Nuclear gene polymorphism (H3), genetic diversity and demographic analyses in *Callinina georgiana*. *N =* number of sequenced samples, *S* = number of segregating sites, *h =* haplotype diversity, π = nucleotide diversity. Bolded values are significant (P < 0.01).

| Population | $N$ | $S$ | $h$ | $\pi$ | Fu's $F_s$ | Tajima $D$ |
| --- | --- | --- | --- | --- | --- | --- |
| Nor-1 | 7 | 14 | 1.000 | 0.020 | 0.723 | 0.734 |
| Siss-1 | 7 | 30 | 0.857 | 0.046 | 1.072 | 0.543 |
| Hann-1 | 10 | 42 | 0.867 | 0.031 | **-2.544** | **-2.017** |
| Pots-1 | 6 | 10 | 0.800 | 0.017 | 1.685 | 1.308 |
| Pots-2 | 7 | 4 | 0.857 | 0.004 | -0.876 | -0.876 |
| Col-1 | 8 | 44 | 0.964 | 0.043 | -1.892 | **-1.668** |
| SCol-1 | 8 | 9 | 0.893 | 0.013 | 0.844 | 0.847 |
| Stark-1 | 10 | 7 | 0.867 | 0.006 | -1.266 | -1.116 |
| Eel-1 | 9 | 17 | 0.944 | 0.017 | -1.349 | -1.162 |
| Cant-1 | 10 | 13 | 0.933 | 0.011 | -1.317 | -1.249 |
| Mad-1 | 9 | 8 | 0.917 | 0.007 | -1.261 | -1.026 |
| Geo-1 | 8 | 39 | 0.893 | 0.049 | 0.329 | -0.238 |
| Ont-1 | 10 | 21 | 0.956 | 0.020 | -0.857 | -0.969 |
| Pau-1 | 8 | 39 | 0.964 | 0.005 | 0.348 | -0.166 |
| Whe-1 | 12 | 2 | 0.409 | 0.002 | 1.017 | 0.687 |
| Cyp-1 | 12 | 0 | 0.000 | 0.000 | -0.820 | -1.215 |

## 4. DISCUSSION

*Callinina georgiana* is arguably the most invasive aquatic invertebrate in the Adirondack region of northern New York. Interestingly, the species was introduced into the region during the mid 1800s – which coincided with when the first dams were being constructed on the Raquette River. The earliest report of an introduction of *C. georgiana* in the northeast was an intentional release of several hundred snails into the Hudson River drainage system (Erie Canal and Mohawk River) by an amateur conchologist in 1867 (Jokinen, 1992; Mills et al., 1993). A few decades later in 1914 the snail was reported in Lake Erie and then in Lake Michigan by 1926. This study is the first attempt at elucidating the population genetic patterns of *C. georgiana*.

Our study revealed that while both the COI and H3 markers exhibited comparable levels of polymorphism, their connectivity patterns across the sampling sites were incongruent with regards to *C. georgina’*s native range. Specifically, the COI marker indicated high levels of genetic connectivity in the snail’s invasive range but there was a marked divergence of haplotypes for native and invasive populations. Conversely, the H3 marker showed a more panmictic-like pattern, with the dominant haplotype being prevalent in both native and invasive populations. This observed mitonuclear discordance could be due to several factors. Given that nuclear genes generally evolve slower than mitochondrial genes, leading to the retention of ancestral genetic variation over longer time periods, the discordance might reflect incomplete lineage sorting, which has been reported in other animals at both the genetic and genomic levels (Toews & Brelsford, 2012; DeRaad et al., 2023). In this case, the clear differentiation in mtDNA could be reflecting a more recent divergence or selective sweep while the nuclear marker may be retaining a signal of ancestral polymorphism. For example, recent study by Lee et al. (2021) on the mussel, *Mytilisepta virgata* showed that it exhibited marked mtDNA divergence into two distinct geographic lineages which corresponded to different water temperature zones in the northwest Pacific. However, this pattern was not mirrored in their nuclear marker (ITS-1) which showed genetic admixture of both mitochondrial lineages. The initial release of approximately 200 banded mystery snails into the Erie Canal and Mohawk River in 1867 marks a significant event in the species’ history in the region (Lewis, 1872; Jokinen, 1992; Mills et al., 1993; David et al., 2017). Given the relatively short time since this introduction, it is plausible that the H3 marker has not had sufficient time for haplotype divergence to emerge between the invasive populations and their native cohorts in the southern US.

Regarding the dispersal dynamics in the anthropogenically altered Raquette River, our findings suggest that the presence of dams does not significantly impede gene flow. This is evident from the low genetic differentiation (Φ_ST_=0.012) and shared haplotypes between sites like Sissonville and Potsdam, despite being separated by multiple dams. Intriguingly, even populations from isolated water bodies such as Lake George and Paul Smiths shared haplotypes with populations from the main Raquette river, indicating a complex pattern of dispersal. The impact of anthropogenic alterations on river connectivity has been a subject of extensive research, yielding conflicting results. Liu and Hershler (2009) observed that dam construction did not significantly affect gene flow in the endangered Bliss Rapids snail, *Taylorchoncha serpenticola.* This finding is consistent with Ruzich et al. (2019), who reported minimal impact of dams on the genetic diversity of certain fish species. In contrast, Liu et al. (2020) discovered reduced genetic diversity and gene flow in unionid mussel populations separated by dams; a pattern echoed in similar studies on chub (DeHais et al., 2010) and riparian plants (Werth et al., 2014). Barnett et al. (2020) further revealed that impounded crayfish populations (*Faxonius validus* and *Faxonius erichsonianus*) exhibited less genetic diversity and higher differentiation, particularly when impounded for extended periods and located closer to dams. The low genetic differentiation in *T. serpenticola,* as per Liu and Hershler (2009) was attributed to ‘passive dispersal’ during high discharge events, like snowmelt in early spring. This dispersal mechanism, prevalent in freshwater molluscs, involves drifting through dams to downstream areas (Martin et al., 2020). Given that *C. georgiana* produces live young capable of drifting, this could explain the high connectivity in the Adirondacks, especially since many of the dams in this region are small likely allow for significant overflow, thereby facilitating connectivity between reservoirs.

Overall *C. georgiana* exhibited high haplotype and low nucleotide diversity for both markers in this study. Combined with the results from the neutrality tests (negative Fu’s Fs and Tajima’D values), these results are indicative of population expansion, most likely due to multiple bottlenecks along with diversification involving frequent interpopulation dispersal (Holland & Cowie, 2007). This data was further corroborated by the high number of private haplotypes recovered. It was once thought that introduced species should theoretically exhibit low genetic diversity in lieu of the founder effect. However, in reality, haplotype diversity can be elevated in an introduced species if propagule pressure is high due to multiple introductions (Roman & Darling, 2007; Rollins et al., 2013; Yang et al., 2018), which we argue is the case here for *C. georgiana* in the Adirondack region. While native *C. georgiana* snails did have a high level of genetic diversity, it was only marginally higher than their invasive cohorts. Comparable results have been shown in other studies on aquatic molluscs. For example, Dupont et al. (2003) found that *Crepidula fornicata,* an introduced snail in France, exhibited high levels of genetic diversity and behaved genetically and demographically similar to its native cohorts from the United States. A major caveat for this study is that the native snails were sampled from only two populations. Populations drawn from additional population across a broader swatch in the southern United States may reveal a different genetic diversity estimate and possibly some level of gene-flow with invasive populations for the COI marker.

Isolated populations with low levels of genetic diversity are considered prime targets for conservation when dealing with native species, since they are often on the brink of extirpation (Miller et al., 2011). Conversely, for invasive species, such populations can be targeted for eradication as they provide a scientifically informed point for concentrating money and resources. However, the fact that neither dams nor distance significantly impacted the genetic diversity nor connectivity of *C. georgiana* populations in its invasive range means that management of the species at the population level may prove to be difficult. This problem is exacerbated by the fact that the culprit responsible for the observed patterns of diversity is anthropogenic in origin. One potential management strategy would be increasing the number of boat stewards in New York State waterways. While there are several boat decontamination stations in the regions where *C. georgiana* has been frequently reported, these stations are not always adequately staffed, with a number of staff working on a strictly voluntary basis. It is therefore possible that *C. georgiana,* especially juveniles can go unnoticed at these stations by being entrained in residual algae or mudpacks. Considering that tourism in the Adirondacks have exploded over the last few decades, especially since the COVID-19 pandemic (Siskind, 2019; Cerialo, 2021), the opportunities for dispersal across multiple waterbodies in the region have increased significantly. Like many other aspects of conservation planning, cultivating an informed and educated public is still the most important strategy for lowering biosecurity risks. Outreach posters and flyers at boat ramps which highlight not only the visible impacts of invasive species such as *C. georgiana* but also the hidden threat of facilitating connectivity among these populations, should be publicized to stakeholders.

If vectors such as boat traffic can be controlled then it should be possible to manually remove the snail from relatively isolated waterbodies that are not hydrologically connected to major watersheds, as was successfully done for invasive apple snails (*Pomacea* spp.) in Florida (Bernatis & Warren, 2014). Complete removal is not necessary as a small population in the long term is likely to be extirpated as long as they are not “rescued” genetically by migration. Populations with limited gene flow due to reduced vectors, such as boats, are likely to exhibit lower genetic diversity. This decreased diversity can heighten the negative consequences of inbreeding. As a result, these populations may face significant survival challenges (Frankham, 2015).

## 5. CONCLUSION

This is the first comprehensive population genetic study carried out for the banded mystery snail, and as far as we know, is also the first for any of the so-called invasive ‘mystery’ snails. Our results provide new insights into the dispersal of this ovoviviparous snail along multiple riverine systems that have been extensively modified through the creation of dams. In summary, our results found a moderate level of genetic structuring across sampling sites with mitonuclear discordance, possibly due to incomplete lineage sorting. While some structuring was observed in the invasive ranges, overall, dams do not appear to impede gene flow of *C. georgiana* in its invasive range. In our main study area, where *C. georgiana* is invasive, we conclude that anthropogenic-mediated dispersal is most likely shuttling migrants across the New York Great Lakes Basin, resulting in high propagule pressure and a highly connected metapopulation of *C. georgiana.* While other vectors, both natural and anthropogenic, may also be at work, further studies using more high-resolution markers (e.g., microsatellites or RAD-Seq generated SNPs) along with a thorough investigation of those vectors will be needed to confirm this.

## AUTHOR CONTRIBUTIONS

Conceptualisation: AD, WANUA. Developing Methods: AD, WANUA. Conducting the research: WANU. Data analysis: AD, WANAU. Data interpretation: AD, WANUA. Preparation figures & tables: WANU. Writing: AD, WANAU.

## ACKNOWLEDGEMENTS

We would also like to thank the following persons for their invaluable help with sampling: Dr. Jennifer Davinack, Dr. Sampath Weerasinghe, Rose Cotto, Geligne Franklin, Dr. Arthur Bogan, Tom Cooper, Steve Johnson, Jeffrey Garner at the Alabama Department of Conservation and Natural Resources, The Wild Center at Tupper Lake and Jakes by the Water (Hannawa Falls, NY). We would also like to thank Drs. Tom Langen, Susan Bailey and Alan Christian for their valuable input in the planning stages of this project

## FUNDING

Funding for this study was provided by the Great Lakes Research Consortium Small Grants Award.

## CONFLICT OF INTEREST STATEMENT

None

## DATA AVAILABILITY STATEMENT

All sequence data generated from this study is available on NCBI GenBank database. R scripts used for analyses are available on request.

## Notes

### Competing Interest Statement

The authors have declared no competing interest.

