## Supplemental Table 1 for "Genetic diversity and connectivity of the invasive gastropod, *Callinina georgiana* (Caenogastropoda: Viviparidae) across a fragmented riverscape: a mitonuclear perspective"

**Supplementary Table 1**

| **Sampling site** | **Population** | **Lat/Lon** | **Habitat** | **pH** | **T (C)** | **Other notes** |
| --- | --- | --- | --- | --- | --- | --- |
| 1 | *Norwood, NY | 44.751659, -75.005745 | Raquette River | 7.7 | 30 | Closer to Norwood hydroelectric dam , rocky area, *Callinina. georgiana* snails attached to the rocks |
| 2 | *Sissonville, NY | 44.692211, -75.008335 | Raquette River | 7.8 | 28 | Closer to Sissonville hydroelectric dam, lots of shells, live *Callinina georgiana* are near Lotus plants |
| 3 | *Potsdam (site 1, NY | 44.666694, -74.989775 | Raquette River | 8.5 | 23 | Two sites of Potsdam are separated from Potsdam East and Potsdam West hydroelectric dams, collected from upper dam area , *Heterogen japonica* detected with *Callinina georgiana* |
| 4 | *Potsdam (site 2), NY | 44.66838, -74.9897 | Raquette River | 8.1 | 21 | Two sites of Potsdam are separated from Potsdam East and Potsdam West hydroelectric dams, collected from just below the dam. Only *Callinina georgiana* detected |
| 5 | *Hannawa Falls, NY | 44.604994, -74.976662 | Raquette River | 7.5 | 23 | Separated from the Hannawa Falls hydroelectric dam, collected from a boat launching site Only *Callinina georgiana* detected |
| 6 | *Colton, NY | 44.545411, -74.936492 | Raquette River | 7.2 | 22 | Between Higley and Colton hydroelectric dams , collected from boat launching area, muddy water, found in leaf litter, *Heterogen japonica* detected with *Callinina georgiana* |
| 7 | *South Colton Reservoir, NY | 44.524725,-74.86278056 | Raquette River | 7.5 | 23 | Closer to South Colton hydroelectric dam, *Heterogen japonica* detected with *Callinina georgiana* |
| 8 | *Stark Falls Reservoir, NY | 44.451005, -74.766048 | Raquette River | 7.9 | 27 | Closer to Stark hydroelectric dam, rocky area. *Heterogen japonica* detected with *Callinina georgiana* |
| 9 | Raymondville Road | 44.728122,-74.992014 | Raquette River | 7.8 | 32 | Closer to Raymondville hydroelectric dam, not included in analysis since only one *Callinina georgiana*  live snail was found |
| 10 | Yalevillie road | 44.7678, -75.002146 | Raquette River | 7.3 | 29 | Closer to Yaleville hydroelectric dam, not included in analysis since only three live *Callinina georgiana*  snails was found |
| 11 | Parishville | 44.6336111,-74.9738889 | Raquette River | N/A | N/A | Between Hannawa and Potsdam dams, no populations recorded, *Campeloma decisum* detected |
| 12 | Carry Falls | 44.4336111, -74.7483333 | Raquette River | N/A | N/A | Near a boat launch, close to Carry falls Dam, No invasive populations detected. |
| 13 | Tupper Lake | 44.2308333,-74.4647222 | Raquette River | N/A | N/A | Near public boat launch area, no invasive populations recorded, *Campeloma decisum* detected |
| 14 | Wild Center | 44.2183333, -74.4327778 | Raquette River | N/A | N/A | Near a boat launch, natural Oxbow shape in the river, *Campeloma decisum* detected |
| 15 | Higley Falls | 44.5122222, -74.9097222 | Raquette River | N/A | N/A | Near Higley Falls State park, between Colton and South Colton hydroelectric dams, no invasive populations recorded, *Campeloma decisum* detected |
| 16 | Norfolk | 44.8025,-74.9905556 | Raquette River | N/A | N/A | Near Norfolk hydroelectric dam, no invasive populations recorded |
| 17 | Massena 1 | 44.9180556, -74.8808333 | Raquette River | N/A | N/A | Near a bridge area, no invasive populations recorded |
| 18 | Massena 2 | 44.915935, -74.891733 | Raquette River | N/A | N/A | Massena Springs Park, near boat launch, no invasive populations recorded, shells of *Campeloma decisum* recorded |
| 19 | Akwesasne | 44.9830556, -74.6975000 | Raquette River | N/A | N/A | Near the opening to the St. Lawrence River, no invasive populations recorded |
| 20 | Blue mountain Lake | 43.8577778,-74.4316667 | Raquette River | N/A | N/A | Near small boat launch area, no invasive populations recorded |
| 21 | *Eel Weir Park | 44.635437, -75.481232 | Oswegatchie River | 9 | 29 | Collected from a boat launching site  Only *Callinina* *georgiana* detected |
| 22 | *Canton Park | 44.600237, -75.177795 | Grasse River | 8.5 | 22 | Collected from a public park area, *Heterogen japonica* detected with *Callinina georgiana* |
| 23 | *Madrid | 44.748361, -75.13054 | Grasse River | 8.6 | 22 | Collected near a boat launch, South from the Grasse River flood risk reduction dam. Only *Callinina. georgiana* detected. |
| 24 | *Sodus Point | 43.270622, -76.977505 | Lake Ontario | N/A | N/A | Collected by a citizen near the beach area Only *Callinina georgiana* collected. |
| 25 | *Lake George, | 43.494029, -73.676284 | Lake George | 8.7 | 26 | Collected from the Lake George beach area with the aid of a recreational diving crowd. *Heterogen japonica* detected with *Callinina georgiana.* Lots of shells detected from both species. |
| 26 | *Paul Smiths | 44.434166,-74.253888 | Lower St. Regis Lake | 8.5 | 29 | Isolated water body, Collected from the beach area, Only *Callinina georgiana* detected. |
| 27 | *Cypress Creek | 34.934860, -87.762810 | Tennessee River tributary | N/A | N/A | N/A |
| 28 | *Wheeler Creek | 34.654210, -87.252700 | Tennessee River tributary | N/A | N/A | N/A |

* denotes the samples used in the analysis , note that temperatures are recorded in different dates in different summer months , N/A -when data not recorded
