## Supplemental Table 2 for "Genetic diversity and connectivity of the invasive gastropod, *Callinina georgiana* (Caenogastropoda: Viviparidae) across a fragmented riverscape: a mitonuclear perspective"

**Supplementary Table 2- Description of the dams near sampling sites**

| Name of the dam | Owner type | Primary purpose | Nearest Town | Maximum Storage (acre ft) | Height (ft) |
| --- | --- | --- | --- | --- | --- |
| Norwood | Private | Hydro electrical | Norwood | 1736 | 24 |
| Sissonville | Private | Hydro electrical | Hewittville | 205 | 16 |
| Potsdam East | Local government | Hydro electrical | Sissonville | 750 | 7 |
| Potsdam West | Local government | Hydro electrical | Potsdam | 750 | 11 |
| Hannawa | Private | Hydro electrical | Potsdam | 690 | 34 |
| Colton | Private | Hydro electrical | Colton | 620 | 29 |
| Higley | Private | Hydro electrical | Colton | 4446 | 37 |
| South Colton | Private | Hydro electrical | South Colton | 3000 | 42 |
| Stark Falls | Private | Hydro electrical | Stark | 12854 | 17 |
